# The Proposed Bone Post-Arterial Type R Capillaries Resolve into Venous and Fatty Acid-Handling Endothelial Cells

**DOI:** 10.64898/2026.07.27.741009

**Authors:** Sanyam Jain, Jessy Davina Joseph, Hanyu Liu, Nainita Roy, Yang Yang, Siddharth Nishant Mundapat, Radek Machan, Antonino Glaviano, Abhishek Chandra, Rinkoo Dalan, Junyu Chen, Navin Kumar Verma, Martine Cohen-Solal, Makarand V. Risbud, Aline Bozec, Anjali P. Kusumbe

## Abstract

Endothelial specialization is increasingly recognized as a fundamental regulator of tissue homeostasis, yet the cellular diversity of the skeletal vasculature remains incompletely resolved. Here, we integrate large-scale single-cell transcriptomics, cross-tissue comparisons, and imaging to comprehensively define endothelial heterogeneity across the skeleton. Our analyses demonstrate that the proposed post-arterial “type R” endothelial population is not a distinct endothelial subtype but instead comprises canonical venous endothelial cells and fatty acid-handling endothelial state. RNA velocity supports a venous continuum, while the proposed type R markers FMO2, and AQP7 lack both endothelial and skeletal specificity. The fatty acid-handling endothelial state, characterized by Lpl and Cd36 is conserved across multiple skeletal sites and non-skeletal tissues, indicating a general endothelial metabolic programme. Within bone, this endothelial state expands following high-fat diet and is suppressed during injury. Together, these findings redefine skeletal endothelial heterogeneity and establish the proposed type R population as part of a venous continuum.

## Introduction

The skeletal vasculature is a highly specialized and dynamic organ system that coordinates bone formation, regeneration, hematopoiesis, immune regulation, and metastatic colonization through specialized vascular niches^1–8^. Beyond supplying oxygen and nutrients, endothelial cells actively regulate tissue homeostasis through angiocrine signaling^7,9–14^. Understanding skeletal vascular heterogeneity is therefore crucial because distinct endothelial populations establish specialized niches that differentially regulate bone development, remodeling, regeneration, hematopoiesis, and disease, making accurate endothelial classification fundamental to both basic skeletal biology and therapeutic discovery^15–18^. Advances in genetics, high-resolution imaging, and single-cell transcriptomics have transformed our understanding of bone blood vessels, revealing remarkable endothelial heterogeneity and identifying multiple endothelial populations with specialized functions^19–29^.

The widespread application of single-cell RNA sequencing (scRNA-seq) has accelerated the discovery of putative endothelial subpopulations^30–35^. However, defining new endothelial identities remains challenging because endothelial cells often exist along continuous transcriptional and developmental trajectories rather than discrete cell types^36,37^, while clustering outcomes are strongly influenced by tissue dissociation, sequencing depth, and computational analyses^38–41^. In mineralized tissues such as bone, these challenges are further compounded by technical limitations in imaging, including tissue autofluorescence, limited imaging depth, and incomplete antibody penetration, making independent spatial validation difficult^2,42–44^. An additional challenge is the frequent disconnect between transcriptomic and imaging-based definitions of endothelial identity. Candidate populations identified by scRNA-seq defined by molecular markers that are not subsequently validated *in situ*, whereas imaging studies rely on different canonical endothelial markers to infer the same populations^45,46^. This issue is exemplified by the recently proposed type R endothelium, a putative post-arterial endothelial subtype implicated in skeletal remodeling^45^. Because endothelial classification forms the foundation for mechanistic studies of bone biology and the development of vascular-targeted regenerative therapies, rigorous validation of newly proposed endothelial populations is essential. Here, we systematically re-evaluated the proposed type R endothelium by integrating large-scale single-cell transcriptomic analyses, RNA velocity, cross-tissue comparisons, and high-resolution spatial imaging across multiple independent datasets and skeletal tissues. Our integrated single cell and imaging analyses demonstrate that the proposed type R population comprises canonical venous endothelial cells together with a previously unrecognized fatty acid-handling endothelial population positioned along a continuous venous endothelial trajectory (**Supplementary Fig. 1**). More broadly, our findings refine the current framework of skeletal endothelial heterogeneity and establish a rigorous approach for defining endothelial cell identities by integrating transcriptomic, spatial, and protein-level evidence.

## Results

### Single-Cell Analyses Reveal That a Proposed type R Endothelial Cell Subcluster Represents Venous Endothelial Cells in Bone

Single-cell RNA sequencing (scRNA-seq) offers a powerful approach to resolve cellular heterogeneity within the bone endothelium. To re-examine the proposed type R endothelial population, we reanalyzed publicly available datasets from Mohanakrishnan et al., which profiled bone endothelial cells (**Fig. 1a**). Across these datasets, cells expressing the defining type R markers, *Fmo2*, *Aqp7*, and *Dach1*, were consistently rare and, in several cases, undetectable using the analytical parameters reported in their study (**Supplementary Fig. 2a-m**). In datasets examining parathyroid hormone (PTH)-induced bone remodeling, where expansion of type R endothelial cells was previously reported, we observed no increase in cells expressing the type R signature (**Fig. 1b, Supplementary Fig. 3a**). Similarly, no expansion was detected in other remodeling contexts, including calcitriol treatment and fracture injury (**Supplementary Fig. 2a, e, Supplementary Fig. 3b**). By contrast, canonical arterial endothelial cells exhibited the expected increase following PTH treatment, consistent with previous reports. Notably, cells expressing the type R signature were reduced following PTH treatment (**Fig. 1b, Supplementary Fig. 3a**).

**Fig 1:**
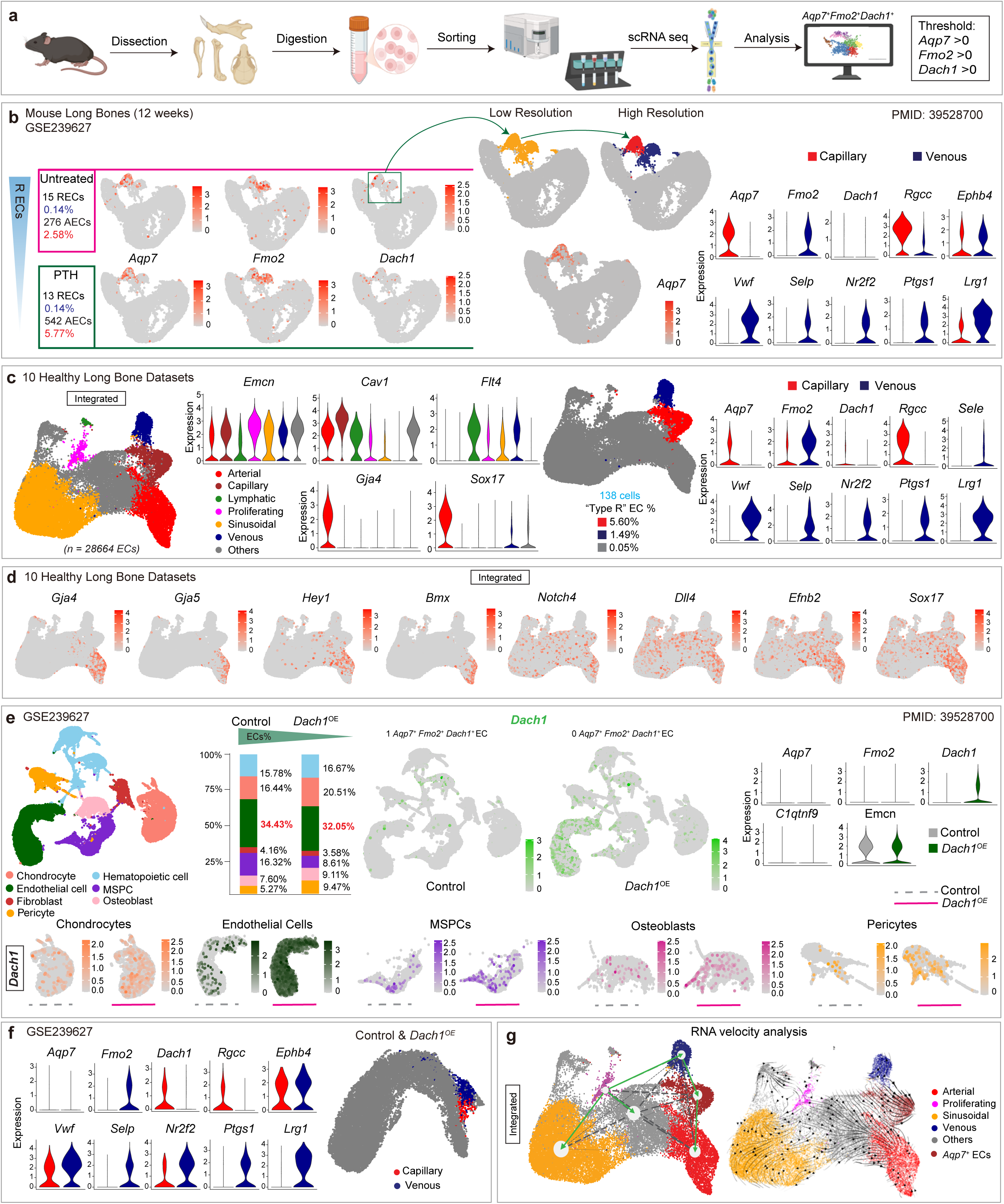
Reassessment of bone endothelial single-cell RNA-sequencing datasets reveals the venous identity of the proposed arterial type R endothelial cluster. a, Schematic representing scRNA-seq experimental workflow and bioinformatic analysis. The expression threshold for markers defining “type R” endothelial state is zero. b, scRNA-seq dataset, GSE239627, profiling sorted endothelial cells (ECs) from young mouse long bones. Feature plots showing expression of *Aqp7, Fmo2* and *Dach1* in endothelial cells between untreated and parathyroid hormone (PTH)-treated group. Fraction of “type R” (*Aqp7^+^Fmo2^+^Dach1^+^*) and arterial (*Cav1^+^Bmx^+^Sox17^+^*) ECs in both the groups is indicated. Resolution of so-called “type R” endothelial cluster into capillary and venous endothelial populations. Violin plots showing expression of *Aqp7, Fmo2, Dach1, Ptgs1, Vwf, Selp, Nr2f2,* and *Lrg1* in capillary (red) and venous (blue) ECs. c, Integrated analysis of ECs from 10 scRNA-seq datasets of healthy young mouse long bones profiling sorted ECs or stromal cells. UMAP showing different endothelial subpopulations. Violin plots showing expression *of Emcn, Flt4, Cav1, Gja4* and *Sox17* across endothelial subtypes. Fraction of “type R” ECs (with respect to total EC population) in capillary, venous and other ECs is indicated. Violin plots showing expression of *Aqp7, Fmo2 and Dach1, Ptgs1, Vwf, Selp, Nr2f2* and *Lrg1* in capillary and venous ECs. d, Feature plots showing expression of *Gja4, Gja5, Hey1, Bmx, Notch4, Dll4, Efnb2, and Sox17* in all ECs from integrated data. e, scRNA-seq dataset, GSE239627, profiling stromal cells from young mouse long bones of control and *Dach1* overexpressing (*Dach1OE*) groups. UMAP projection showing various cell types of bone and bone marrow microenvironment. Stacked bar plots showing the proportion of different cell types in control and *Dach1^OE^* samples. Feature plots showing *Dach1* expression in the control and *Dach1^OE^* samples across different cell types. Violin plots showing expression of *Aqp7, Fmo2, Dach1, C1qtnf9 and Emcn* in control and *Dach1^OE^* samples. f, Combined UMAP showing capillary and venous ECs in control and Dach1 overexpressing (*Dach1^OE^*) samples. Violin plots showing expression of *Aqp7, Dach1, Vwf, Lrg1, Selp, Ptgs1, Nr2f2, Fmo2* in capillary and venous ECs. g, RNA velocity analysis on integrated data of GSE239627, GSE128423, GSE228995, GSE154247, GSE122465, GSE220836, GSE281311 and GSE225429 showing the predicted transcriptional dynamics and lineage relationships among EC subpopulations.

We next evaluated marker specificity and cluster coherence. *Fmo2* and *Aqp7* were each expressed in only approximately 50% of the cell population within the originally annotated type R cluster consistently in multiple clustering attempts and displayed incomplete pairwise co-expression, indicating weak internal consistency of the proposed signature (**Fig. 1b, Supplementary Fig. 3a**). Moreover, these genes were not restricted to a discrete endothelial subpopulation but were variably expressed across other endothelial subsets. To resolve the identity of this population, we performed unbiased reclustering of endothelial cells. This analysis resolved the originally designated “type R” cluster into two transcriptionally distinct endothelial subpopulations (**Fig. 1b**), consistent with the differential expression of *Fmo2* and *Aqp7*, which marked separate subsets within the original cluster. Strikingly, the *Aqp7*- negative subpopulation exhibited robust expression of canonical venous endothelial markers, including *Vwf, Selp,* and *Nr2f2* (**Fig. 1b, Supplementary Fig. 3a**).

To test the robustness of these observations, we integrated and analysed ten published scRNA-seq datasets of sorted endothelial or stromal cells from healthy mouse long bones (**Fig. 1c**, **Table 1**). We used standardized and fully reproducible analytical pipelines, the codes for which are provided in the **Supplementary Information**. Across this integrated atlas of endothelial cells, cells expressing the proposed type R marker signature (*Dach1, Fmo2,* and *Aqp7*) remained extremely rare (**Figure 1c, Supplementary Fig. 3c**). Instead, a substantial proportion of cells assigned to the originally defined type R cluster, here seen as *Emcn^+^ Cav1^+^ Flt4^-^* clusters, displayed a canonical venous endothelial transcriptional program characterized by robust expression of *Vwf, Selp, Sele,* and *Nr2f2* (**Fig. 1c**). Furthermore, canonical arterial markers, including *Gja4, Gja5, Bmx, Dll4, Notch4* and *Hey1*, were not expressed in the proposed type R cluster despite its reported identity as a post-arterial capillary population (**Fig. 1d**). These findings indicate that a subset of the proposed type R population is more consistent with venous endothelium than with a novel post-arterial capillary subtype. This observation also holds true for scRNA-seq dataset obtained from juvenile (P21) mice (**Supplementary Fig. 4a**). We next examined functional validation datasets generated using a *Dach1* overexpression mouse model. We could not detect any increase in type R-associated markers (*Fmo2* or *Aqp7*), but reduced endothelial cell abundance (**Fig. 1e, f, Supplementary Fig. 4b, c**). These findings contrast with the previous report proposing DACH1 as a positive regulator of type R-associated transcriptional programs and further underscore the inconsistency in defining this population.

To further investigate developmental relationships among endothelial subpopulations, we performed RNA velocity analysis^47^, a computational approach that infers future transcriptional states by quantifying the relative abundance of unspliced and spliced mRNA transcripts (**Fig. 1g**). Unlike conventional clustering, which provides a static snapshot of cellular identities, RNA velocity reconstructs dynamic cell-state transitions and enables inference of potential lineage trajectories within heterogeneous populations. RNA velocity analysis revealed a continuous transcriptional relationship between the venous endothelial cell cluster and the *Aqp7*-positive endothelial cluster rather than supporting the existence of a discrete post-arterial endothelial lineage (**Fig. 1g**). The inferred velocity vectors consistently connected venous endothelial cells to both arterial endothelial cells and the *Vwf*-negative, *Aqp7*-positive population (**Fig. 1g**), suggesting that these populations occupy related positions along a shared endothelial differentiation continuum. Importantly, the directionality of the velocity streams indicated that venous endothelial cells represent an upstream cellular state capable of giving rise to both arterial endothelial cells and the *Aqp7*-positive subset. This observation is in line with previous studies showing venous endothelial cells expand the vascular bed giving rise to arterial endothelial cells^48–51^. Collectively, RNA velocity analysis provides an independent and orthogonal line of evidence that at least one subset of the proposed type R population corresponds to venous endothelium, while the other occupies an intermediate position within the broader venous-to-arterial differentiation continuum rather than representing a distinct post-arterial subtype. Interestingly, both clusters were not only present in bone and bone marrow but also in the periosteum, where the *Aqp7*-positive cluster constituted the majority of endothelial cells (**Fig. 2a, b**). Imaging analyses also confirm the presence of vWF-expressing veins (**Fig. 2c**).

**Fig 2:**
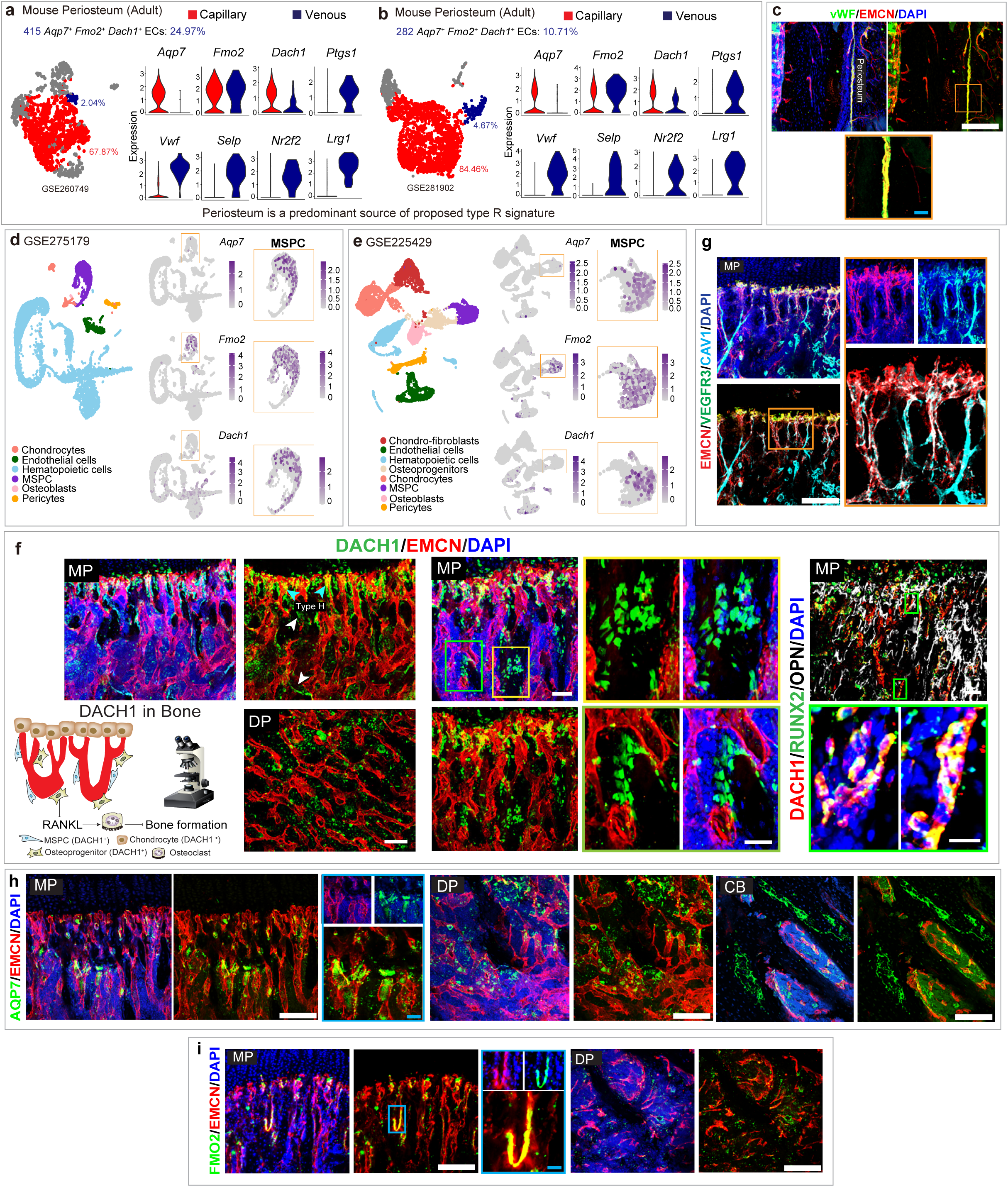
Proposed type R markers are not endothelial-specific and DACH1 protein is predominantly expressed in osteoprogenitors and type H vessels. **a**, scRNA-seq dataset, GSE260749, profiling periosteal cells from adult mice. UMAP showing capillary and the venous ECs. Fraction of “type R” ECs is mentioned. Violin plots showing the expression of *Aqp7, Fmo2, Dach1, Ptgs1, Vwf, Selp, Nr2f2* and *Lrg1* in capillary and venous ECs. **b**, scRNA-seq dataset, GSE281902, profiling periosteal cells from adult mice. UMAP showing the capillary and the venous ECs and the proportions of “type R” ECs. Violin plots showing the expression of *Aqp7, Fmo2, Dach1, Ptgs1, Vwf, Selp, Nr2f2* and *Lrg1* in the capillary and the venous ECs. **c**, Representative images of periosteum of young murine femurs showing immunolabelling for VWF (green) and Endomucin (red) nuclear staining with DAPI (blue). Scale bars: 70 μm (white) and 10 μm (cyan). **d**, scRNA-seq dataset, GSE275179, profiling stromal cells from young murine long bones. Feature plots showing expression of *Aqp7*, *Fmo2* and Dach1 across all cells. Note the expression of *Aqp7, Fmo2* and *Dach1* in MSPCs. **e**, scRNA-seq dataset, GSE225429, profiling stromal cells from young murine long bones. Feature plots showing expression of *Aqp7*, *Fmo2* and *Dach1* across all cells. Note the expression of *Aqp7, Fmo2* and *Dach1* in MSPCs. **f**, Representative images of the metaphyseal (MP) and diaphyseal (DP) region of young murine tibia immunolabelled for DACH1 (green) and Endomucin (red) and nuclear stained with DAPI (blue). Representative images of MP regions of young murine tibia showing immunolabelling for DACH1 (red), RUNX2 (green) and Osteopontin (white) and nuclear staining with DAPI (blue). Scale bars: 50 μm (white). **g**, Representative images of MP region of young murine tibia showing immunolabelling for VEGFR3 (green), Endomucin (red) and Caveolin-1 (cyan) nuclear staining with DAPI (blue). Scale bars: 70 μm (white) and 10 μm (cyan). **h**, Representative images of MP, DP regions and cortical bone of young murine tibia showing immunolabelling for AQP7 (green), Endomucin (red) and nuclear staining with DAPI (blue). Scale bars: 70 μm (white) and 10 μm (cyan). **i**, Representative images of MP and DP regions of young murine tibia showing immunolabeling for FMO2 (green) and Endomucin (red) and nuclear staining DAPI (Blue). Scale bars: 70 μm (white) and 10 μm (cyan).

### Genetic and functional analyses do not support an endothelial-specific role for *Dach1* in bone remodelling

Analysis of individual datasets revealed that cells expressing the proposed type R markers (*Fmo2*, *Aqp7*, and *Dach1*) were consistently rare across all examined datasets and, in several cases, entirely undetectable (**Fig. 1b, Supplementary Fig. 2a-m**). Although *Dach1* has been proposed as a key transcriptional regulator of type R cells, *Dach1* was not identified as a differentially expressed gene in the original dataset (**Supplementary Fig. 4a**). Reanalysis of the *Dach1* overexpression (*Dach1*- OE) dataset confirmed the absence of type R cell expansion (**Supplementary Fig. 4b, c**). On the contrary, the overall proportion of endothelial cells was reduced in *Dach1*-OE bones compared to controls (**Fig. 1e**). Furthermore, *Dach1* expression was not restricted to endothelial cells but was broadly distributed across bone marrow stromal and haematopoietic compartments, indicating a lack of endothelial specificity (**Fig. 2d, e**).

Consistent with these transcriptomic findings, imaging analyses using the same antibody as in the original study showed that DACH1 protein is predominantly expressed in perivascular, mesenchymal stem/progenitor, and osteolineage cells (**Fig. 2f, Supplementary Fig. 4d**). This pattern aligns with prior studies linking DACH1 to osteoprogenitor function and bone formation^52,53^. In contrast, DACH1 signal in vascular capillaries in the regions of bone, where type R capillaries has been reported, was indistinguishable from background.

DACH1 is readily detectable in osteolineage populations, whereas its expression in capillaries in bone is weak or inconsistent (**Fig. 2f, Supplementary Fig. 4d**). Thus, the phenotypes attributed to endothelial-specific *Dach1* manipulation can be parsimoniously explained by direct effects on osteoprogenitors and osteoblasts, rather than by the activity of a specialized endothelial subset. Collectively, these findings highlight the importance of precise lineage specificity when interpreting Cre-based genetic models in bone. A revised framework is therefore warranted, in which the observed bone anabolic effects are attributed to osteolineage-driven mechanisms rather than endothelial-specific *Dach1* function. Future studies should incorporate more stringent endothelial-restricted genetic strategies to resolve lineage contributions with greater fidelity.

### Integrated Transcriptomic and Imaging Analyses Redefine Bone Endothelial Marker Specificity

To complement the transcriptomic analyses, we systematically evaluated the spatial localization and protein-level expression of markers used to define the proposed type R endothelial population. Previous studies relied heavily on immunostaining for Caveolin and Endomucin to delineate type R capillaries. However, defining markers of the type R cluster identified by scRNA-seq were *Fmo2*, *Aqp7*, and *Dach1*. Notably, Caveolin and Endomucin label heterogeneous endothelial compartments across diverse vascular states (**Fig. 2g**). Consequently, the imaging framework previously used to support the existence of type R vessels lacks spatial specificity. Moreover, it does not incorporate the key markers (*Fmo2*, *Aqp7*, and *Dach1*) identified in scRNA-seq studies, which are essential to substantiate a distinct anatomical or functional identity.

DACH1 expression was not detected in endothelium in the bone regions where type R endothelium has been previously reported (**Supplementary Fig. 4e**). In addition to *Dach1*, *Fmo2* and *Aqp7* have been proposed as defining markers of the type R cluster based on transcriptomic enrichment. However, there is a complete lack of corresponding protein-level validation and spatial imaging for these candidates. This omission is particularly significant because *Dach1*, *Fmo2* and *Aqp7* are not endothelial-specific markers but are broadly expressed in bone and bone marrow cells (**Fig. 2d, e**). Since both Fmo2 and Aqp7 are known for their stress-responsive functions, their expression may reflect transient hemodynamic or physiological states induced during single-cell isolation process rather than stable endothelial identity^54–57^. Both *Fmo2* and *Aqp7* are responsive to shear stress and metabolic cues, suggesting that their expression may reflect transient hemodynamic or physiological states rather than stable endothelial identity. This raises an important technical consideration: transcriptional signatures detected in single-cell RNA sequencing datasets may be influenced by tissue dissociation, enzymatic processing, and flow cytometric isolation, all of which are known to perturb endothelial gene expression profiles. Thus, the apparent enrichment of these genes may represent an artifact of experimental conditions rather than evidence of a bona fide endothelial expression. Most importantly, AQP7 was not detected in bone or bone marrow endothelium while FMO2 was detected only in a small subset of column-like Type H vessels in the metaphysis (**Fig. 2h, i**). Taken together, these imaging and protein-level analyses fail to provide evidence for a spatially or molecularly distinct type R endothelial population. Instead, they reveal that the markers used to define this cluster lack specificity, are broadly distributed across vascular and non-vascular compartments, and in several cases are not expressed in endothelial cells at all. Critically, the absence of spatial restriction, combined with the lack of endothelial-specific protein validation, expression of venous signature rather than proposed arterial phenotype undermines the central premise of a discrete type R vascular identity.

### Type R Markers Are Widely Expressed Across Soft Tissues and Lack Bone- Endothelial Specificity

The absolute lack of spatial or proteomic validation of computationally derived type R markers from transcriptomic data is further compounded by cross-tissue analyses, which demonstrate that *Fmo2*, *Aqp7*, and *Dach1* are not restricted to bone endothelium but are broadly expressed across diverse organs, including the heart, colon, brain, lung, skeletal muscle, and liver (**Fig. 3a**). Notably, triple-positive (*Fmo2⁺Aqp7⁺Dach1⁺*) populations are readily detected in these soft tissues across multiple independent datasets, whereas in bones, they are not (**Fig. 3b**). This indicates that these genes reflect conserved, systemic transcriptional programs rather than a bone-specific endothelial identity. Moreover, *Dach1*, proposed as a key transcriptional regulator of type R endothelial cells, is expressed at even higher levels in endothelial cells from multiple other tissues than in bone, further calling into question its specificity for this population (**Fig. 3b**).

**Fig 3:**
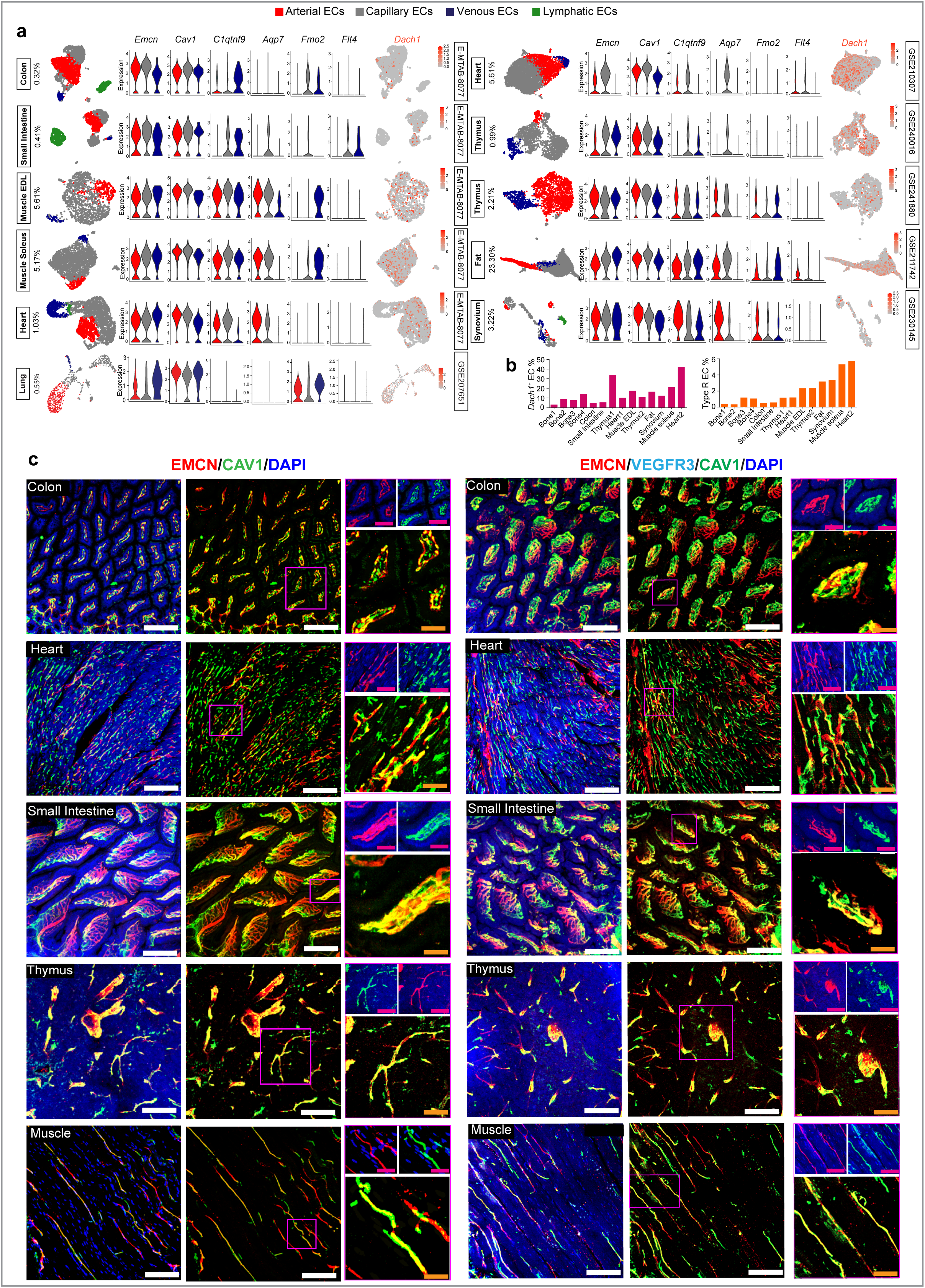
Proposed type R marker expression is not bone-specific. **a**, Vascular analysis of publicly available scRNA-seq datasets of colon, small intestine, muscle, heart, thymus, fat, synovium and lung profiling sorted endothelial cells (ECs) or stromal cells. UMAPs showing arterial, venous, capillary and lymphatic ECs. Fraction of “type R” ECs is mentioned. Violin plots showing expression of *Emcn, Cav1, C1qtnf9, Aqp7, Fmo2,* and *Flt4* and feature plots showing Dach1 expression across endothelial subtypes. **b**, Bar plots indicating the proportion of *Dach1⁺* ECs and “type R” ECs across tissues. **c**, Representative images of murine non-skeletal tissues including colon, heart, small intestine, thymus, and skeletal muscle showing immunolabelling for Endomucin (red), Caveolin-1 (green) and VEGFR3 (cyan) and nuclear staining with DAPI (blue). Scale bars: 70 μm (white), 50 μm (yellow) and 20 μm (magenta).

Within bone, these markers (*Dach1*, *Aqp7*, *Fmo2*) are not exclusive to endothelial cells and are expressed by other stromal and non-endothelial populations, further undermining their specificity for defining a discrete vascular subtype. Analyses of Endomucin and Caveolin-1 expression, which were previously used to identify type R vessels by imaging, similarly demonstrate widespread distribution. Endomucin-positive, Caveolin-1-positive, and VEGFR3-negative capillaries are commonly found across many soft tissues, including the intestine, skeletal muscle, heart, and thymus (**Fig. 3c**).

### Fatty Acid–Handling Endothelial Cells Represent a Component of the Venous Endothelial Continuum

To further resolve the identity of the *Aqp7⁺* endothelial cells previously grouped within the proposed remodeling endothelial cell (REC) cluster, we examined the transcriptional characteristics of the two populations that emerged from high-resolution reclustering analyses. As shown above, one subpopulation displayed a clear transcriptional profile of venous endothelial cells. In contrast, examination of the second population exhibited strong signatures associated with fatty acid transport and lipid handling (**Fig. 4a**). We therefore termed this latter population “endothelial cells (FA-ECs)”.

**Fig 4:**
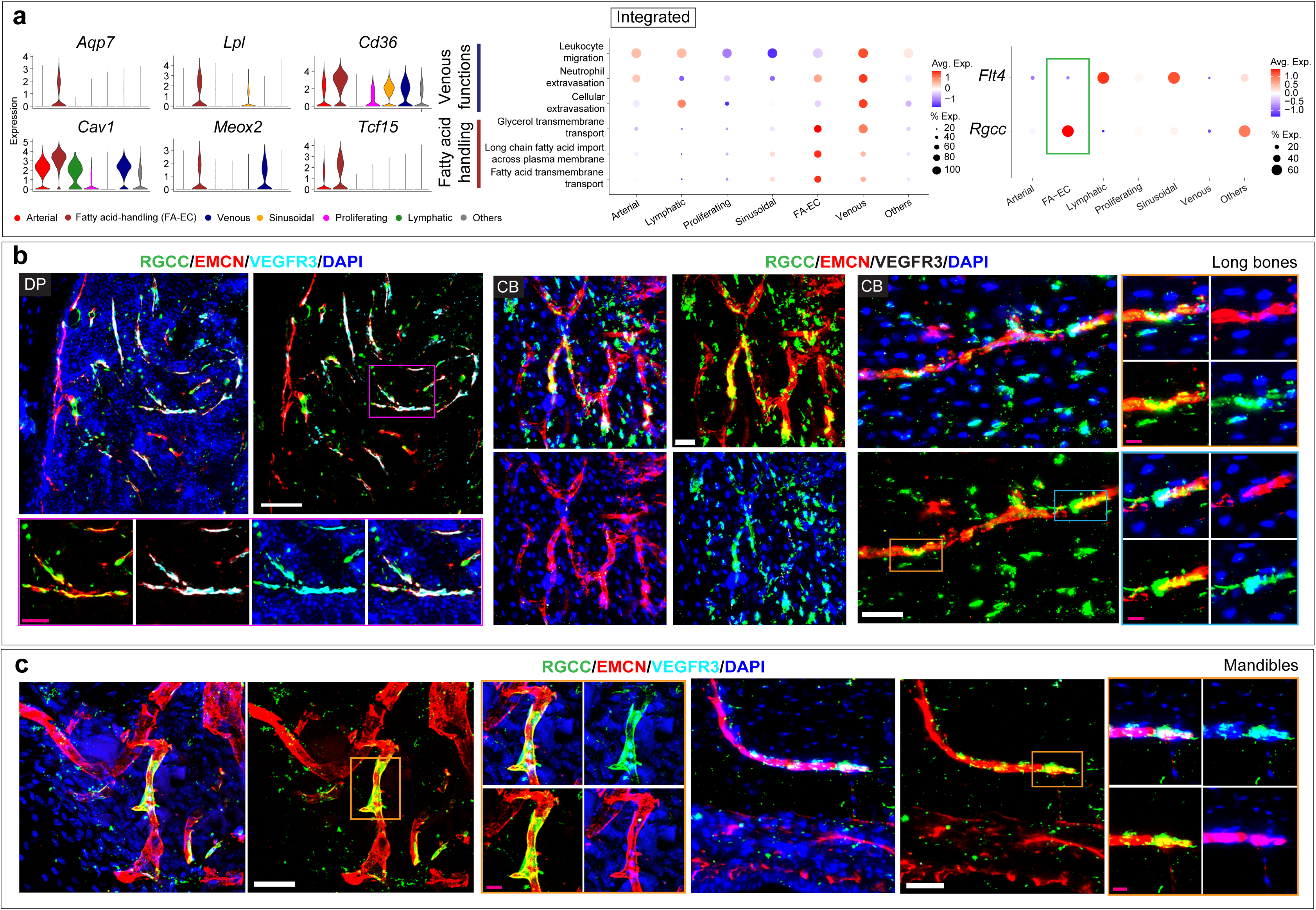
Fatty acid-handling endothelial cells form part of the venous continuum within the proposed type R endothelial cluster. **a**, Violin plots showing expression of *Ap7, Lpl, Cd36, Cav1, Meox2,* and *Tcf15* across endothelial sub-populations from the integrated analysis of 10 healthy long bone scRNA-seq datasets. Dot plot showing scores of venous function and fatty acid handling related gene ontology terms across endothelial sub-populations. Dot plot showing differential expression of *Rgcc* and *Flt4* across endothelial sub-populations in integrated data. **b**, Left panel: Representative images of diaphyseal (DP) region of young murine tibia showing immunolabelling for RGCC (green), Endomucin (red) and VEGFR3 (cyan) and nuclear staining with DAPI (blue). Right panel: Representative images of cortical bone (CB) of young murine tibia showing immunolabelling for RGCC (green), Endomucin (red) and VEGFR3 (white) and nuclear staining with DAPI (blue). **c**, Representative images of young murine mandible showing immunolabelling for RGCC (green), Endomucin (red) and VEGFR3 (cyan) and nuclear staining with DAPI (blue). Scale bars: 50 μm (white) and 15 μm (magenta).

Functional gene ontology analyses support these annotations. Venous endothelial cluster scores high for biological processes associated with leukocyte trafficking and vascular permeability, including neutrophil extravasation, cellular extravasation and neutrophil migration (**Fig. 4a**). In contrast, FA-ECs exhibit upregulation of pathways involved in lipid metabolism, including glycerol transmembrane transport, fatty acid transmembrane transport and long-chain fatty acid import across the plasma membrane (**Fig. 4a**). These transcriptional features closely resemble specialized lipid-handling capillary endothelial populations previously described in adipose tissue, skeletal muscle and cardiac vasculature^30,31,58–61^.

Because unsupervised clustering can artificially partition continuous transcriptional states, we systematically assessed the effect of clustering resolution on endothelial population structure. At lower clustering resolutions, venous endothelial cells and FA- ECs merged into a single endothelial population occupying one end of the endothelial continuum (**Supplementary Fig. 5a**). Furthermore, expression of the proposed REC markers was not uniformly distributed across the two populations. Whereas *Aqp7* expression was largely restricted to FA-ECs, *Fmo2* was preferentially enriched in venous population (**Supplementary Fig. 5b**). Reanalysis of the original dataset used in the study to define RECs confirmed that increasing clustering resolution reproducibly separated the REC cluster into transcriptionally distinct venous and fatty acid-handling endothelial populations, indicating that the REC identity arises from insufficient resolution of a broader endothelial continuum rather than representing a discrete endothelial subtype (**Supplementary Fig. 5c**).

To determine whether FA-ECs are restricted to long bones or represent a conserved endothelial population throughout the skeleton, we reanalyzed scRNA-seq datasets from two craniofacial skeletal sites, the calvarium and mandible. A distinct FA-EC population was identified in both skeletal sites. Although FA-ECs and venous endothelial cells remained transcriptionally related, they exhibited greater separation in craniofacial endothelial landscapes than in long bones, suggesting skeletal site-specific adaptations of endothelial identity (**Supplementary Fig. 6a-c**). Imaging analysis with the markers expressed by FA-ECs further confirmed the presence of FA- ECs in long bones and mandible (**Fig. 4b, c, Supplementary Fig. 6d**). The lipid- association of this FA-EC population is further exemplified by its expansion in response to high fat diet (**Fig. 5a**). Previously, two artery-like endothelial cell (AL-EC) populations were reported in calvarium (PMID: 38834586), which actually correspond to venous and FA-EC identities based on co-expression of proposed AL-EC markers and canonical venous/FA-EC markers in respective clusters (**Fig. 5b**). Further, we assessed the expression patterns of proposed markers of AL-EC populations in our integrated data of long bone endothelial cells. Like calvarium, markers of AL-EC populations 1 and 2 enrich in FA-EC and venous ECs clusters respectively (**Fig. 5c**). To investigate the role of FA-ECs during tissue injury, we analyzed endothelial populations during fracture repair. A marked reduction in FA-EC abundance was observed after fracture in both whole-bone and periosteal preparations (**Fig. 5d, e**). Key components of the FA-EC fatty acid uptake program, including *Lpl*, *Aqp7*, *Cd36*, *Tcf15*, and *Meox2* were significantly downregulated. A similar response was observed in periosteal FA-ECs and calvarial FA-ECs, both of which exhibited a pronounced decline after defect and injury (**Fig. 5f, g**). Together, these observations reveal a conserved endothelial response of FA-ECs to injury across distinct skeletal compartments characterized by repression of fatty acid transport functions.

**Fig 5:**
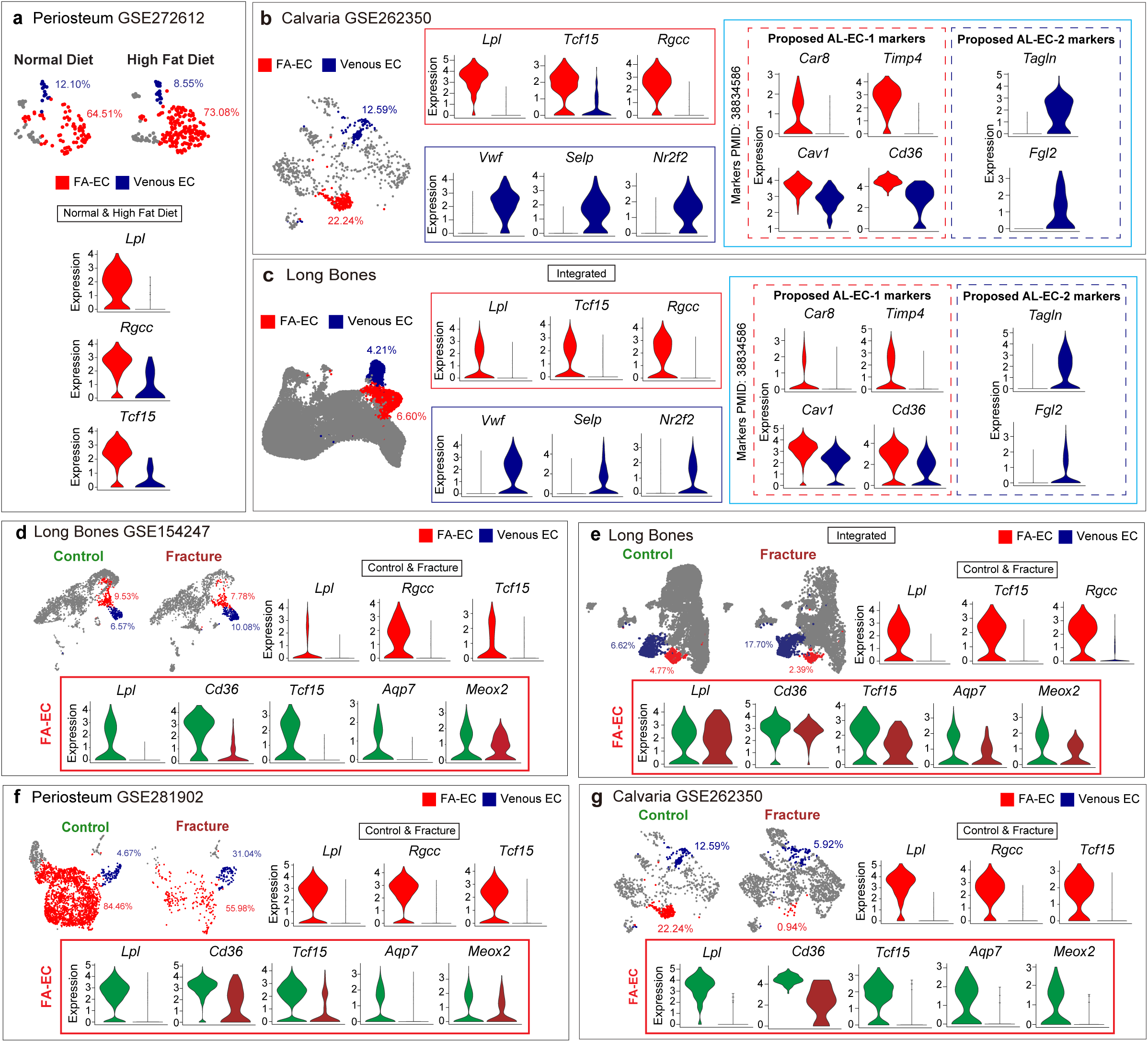
Fatty acid-handling endothelial cells are broadly distributed across the skeletal vasculature. **a**, scRNA-seq dataset, GSE272612, profiling periosteal cells from adult mice fed with normal and high fat diet. UMAPs showing fatty acid-handling endothelial cells (FA- ECs) and venous ECs in periosteum of mice from normal diet vs high fat diet group. **b**, scRNA-seq dataset, GSE262350, profiling stromal cells of young murine calvarium. UMAP showing FA-EC and venous EC clusters in young murine calvarium. Violin plots showing expression of *Lpl, Tcf15, Rgcc, Vwf, Selp, Nr2f2, Car8, Timp4, Cav1, Cd36, Tagln,* and *Fgl2* in FA-ECs vs venous ECs. **c**, UMAP showing FA-EC and venous EC clusters from the integrated analysis of 10 healthy long bone scRNA-seq datasets. Violin plots showing expression of *Lpl, Tcf15, Rgcc, Vwf, Selp, Nr2f2, Car8, Timp4, Cav1, Cd36, Tagln,* and *Fgl2* in FA-ECs vs venous ECs. **d**, UMAPs showing FA-EC and venous EC clusters in control vs fractured (post-fracture day 14) young murine tibia from GSE154247 dataset. Violin plots showing expression *of Lpl, Tcf15, Rgcc* in FA-ECs vs venous ECs of both groups combined. Violin plots showing expression of *Lpl*, *Cd36*, *Tcf15*, *Meox2*, and *Aqp7* in FA-ECs of control vs fracture groups. **e**, UMAPs showing FA-EC and venous EC clusters in control vs fractured young murine tibia from integrated analysis of datasets GSE154247, GSE128423, GSE228995, GSE242712, GSE265817, and GSE266774. Violin plots showing expression of *Lpl, Tcf15, Rgcc* in FA-ECs vs venous ECs of both groups combined. Violin plots showing expression of *Lpl*, *Cd36*, *Tcf15*, *Meox2*, and *Aqp7* in FA-ECs of control vs fracture groups. **f**, UMAPs showing FA-EC and venous EC clusters in periosteum from control vs fractured (post- fracture day 7) young murine bones from GSE281902 dataset. Violin plots showing expression of *Lpl, Tcf15, Rgcc* in FA-ECs vs venous ECs of both groups combined. Violin plots showing expression of *Lpl*, *Cd36*, *Tcf15*, *Meox2*, and *Aqp7* in FA-ECs of control vs fracture groups. **g**, UMAPs showing FA-EC and venous EC clusters in control vs fractured (post-injury day 14) young murine calvarium from GSE262350 dataset. Violin plots showing expression of *Lpl, Tcf15, Rgcc* in FA-ECs vs venous ECs of both groups combined. Violin plots showing expression of *Lpl*, *Cd36*, *Tcf15*, *Meox2*, and *Aqp7* in FA-ECs of control vs fracture groups.

## Discussion

The skeletal vasculature comprises specialized endothelial populations that regulate bone development, remodeling, regeneration, ageing, and disease. Defining these endothelial identities is therefore fundamental to understanding skeletal biology and developing vascular-targeted therapies^62–64^. By integrating large-scale single-cell transcriptomic analyses, RNA velocity, cross-tissue comparisons, genetic datasets, and high-resolution spatial imaging, our study revises the current framework of skeletal endothelial heterogeneity. Rather than supporting a discrete post-arterial type R endothelial subtype, our findings demonstrate that the proposed population comprises canonical venous endothelial cells together with a metabolically specialized fatty acid-handling endothelial population positioned along a continuous venous endothelial trajectory (Fig. 6). Interestingly, periosteal datasets contained a substantially higher proportion of fatty acid-handling endothelial cells (FA-ECs). A major biological finding of this study is the identification of fatty acid-handling endothelial cells (FA-ECs) as a conserved endothelial population throughout the skeleton. Metabolically specialized endothelial cells that regulate fatty acid uptake have previously been described in soft tissues, including the heart, adipose tissue, and skeletal muscle, where they facilitate tissue-specific lipid metabolism^30,31,58–61^. FA- ECs are characterized by transcriptional programmes associated with lipid uptake and fatty acid transport and undergo coordinated suppression during fracture repair across long bone, periosteum, and calvarium, indicating that endothelial metabolic programmes are dynamically remodelled during skeletal regeneration.

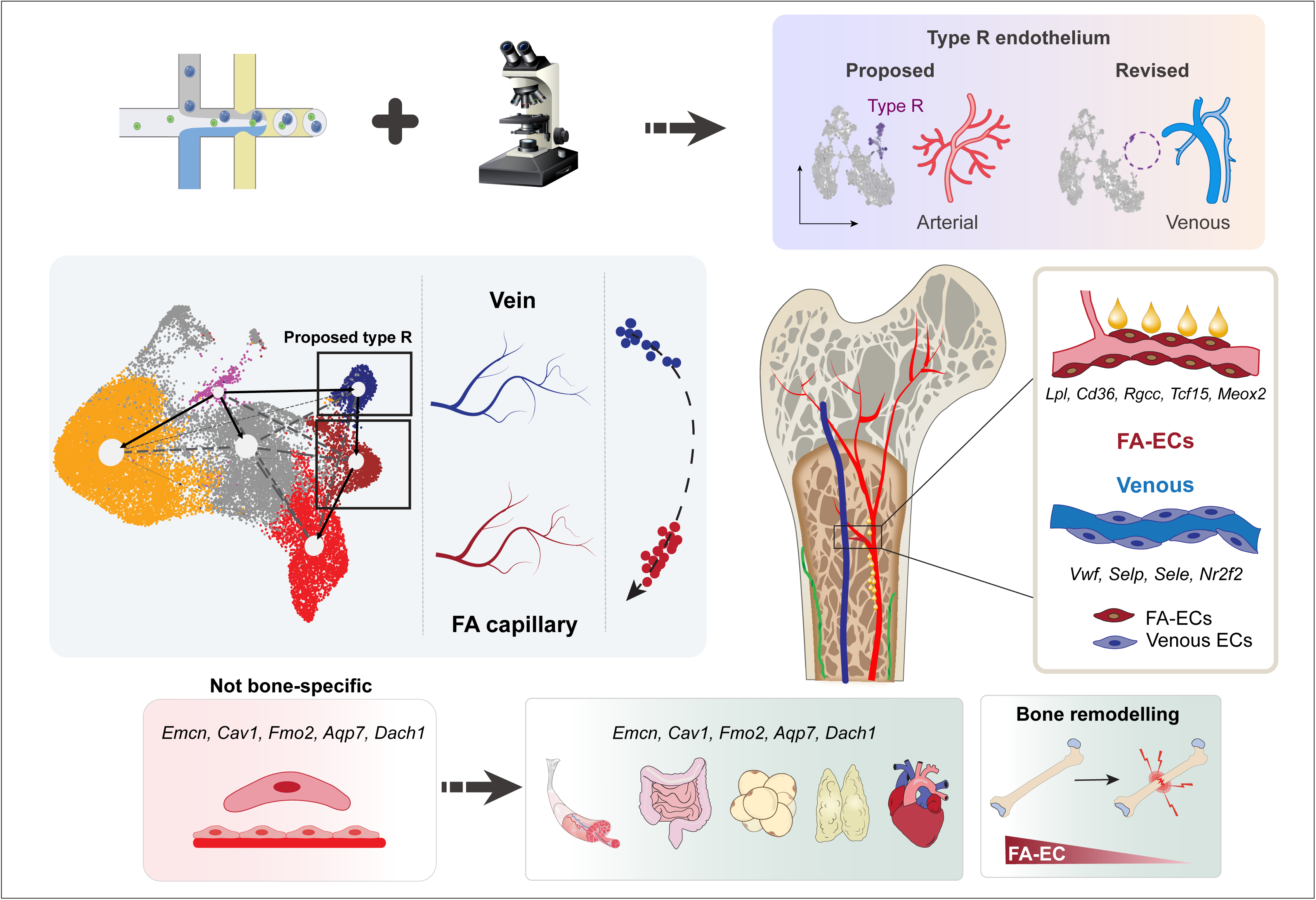

More broadly, our study highlights a fundamental principle for interpreting endothelial heterogeneity from single-cell atlases. Computationally identified transcriptional clusters should not be equated with novel cell types without rigorous orthogonal validation. Consistent with this principle, the proposed murine type R signature also fails to define a distinct endothelial population in human bone datasets, where the corresponding transcriptional programme instead aligns with a fatty acid-handling endothelial phenotype. Furthermore, the proposed type R markers DACH1, FMO2, and AQP7 are neither endothelial- nor bone-specific but are broadly expressed across multiple tissues and cell types. Likewise, the imaging markers used to define type R vessels, including Endomucin and Caveolin-1, are widely expressed in vascular beds of diverse soft tissues, including the heart, skeletal muscle, intestine, thymus, lung, and adipose tissue. Thus, neither the transcriptomic markers nor the imaging criteria previously used to define type R endothelium are unique to skeletal vessels.

Along the same lines, canonical murine type R markers fail to label a clear EC cluster in human scRNA-seq data as well. Using label transfer from mouse to human ECs, type R ECs have been shown in human bone scRNA-seq data^65^, where multiMAP integration results divides proposed type R cluster into two separate islands, however, only one of them is described as type R, which is enriched in *MGLL* (Monoglyceride lipase), *CD36*, *RUNX1T1* (encoding adipogenic regulator CBFA2T1) indicating FA-EC phenotype. Differing marker combinations have been used to define bone endothelial subpopulations. This has resulted in substantial variability in the identification and interpretation of seemingly same cell populations. Striking contradictions emerge when comparing published datasets. For example, in the same study proposing “type R” endothelial population, type H endothelial cells, referred to as mpECs in that study, were reported to increase with age (Figure 8e and Figure 1d of PMID: 39528700)^45^, whereas multiple studies^3,8,13,66^ report a significant age-associated decline in Type H endothelial cells. Similarly, reports of increased arterial endothelial cells in aging (Figure 1d of PMID: 39528700)^45^ bone contrast with previous conclusions from the same group and other studies^1,67^. Angiogenesis and osteogenesis in the skull have been recently reported to be uncoupled and associated with tip cells (PMID: 38834586), which contrasts with earlier work describing Type H vessels in skull bone^68^ lacking tip cells and supporting coordinated angiogenesis–osteogenesis coupling^4,43^. Collectively, our study provides a revised framework for skeletal endothelial biology in which endothelial heterogeneity is organized as a continuum of anatomical and metabolic specialization rather than an expanding catalogue of discrete endothelial subtypes. Beyond redefining the proposed type R population, our findings establish general principles for rigorous endothelial classification that will facilitate future studies of vascular regulation in bone homeostasis, regeneration, ageing, and disease. More broadly, this integrative framework provides a conceptual blueprint for distinguishing stable endothelial identities from dynamic physiological states across vascular beds in multiple organs.

## Resource availability

Further information and requests for resources and reagents should be directed to, and will be fulfilled by, the lead contact, Anjali P. Kusumbe

## Materials availability

This study did not generate new unique reagents.

## Data and Code Availability

All codes used for the analysis of single-cell RNA sequencing data is provided in the Supplementary Information.

## Acknowledgements

The author(s) declare that financial support was received for the research. A. P. K. is supported by Ministry of Education (MOE) Singapore: Academic Research Funds (#024983-00001), European Research Council (StG: metaNiche, 805201), European Union’s Horizon 2020 (No 857524). S. J. is supported by the Nanyang Technological University Research Scholarship (Reg. No. 200604393R). Y.Y. is supported by National Nature Science Foundations of China (No. 824B2026). A. C. is supported by R01 AG082681 grant. R. D. is supported by Personalised Cardiometabolic Risk Management (predict-2-prevent) and the vascular research initiative supported by National Healthcare Group and Lee Kong Chian School of Medicine. J. C. is supported by the National Nature Science Foundations of China (Nos. 82422021, 82270961), and Sichuan Science and Technology Program (No. 2023JDRC0018). N. K. V. acknowledges funding support, in part, by the National Research Foundation Singapore under its Open Fund - Individual Research Grant administered by the Singapore Ministry of Health’s National Medical Research Council (#MOH-001892). M.V.R. acknowledges support by the National Institute on Aging (R01AG073349) and National Institute of Arthritis and Musculoskeletal and Skin Diseases (R01AR055655 and R01AR082460). A.B. received funding from DFG-SFB/TRR369 DIONE- 501752319.

## Methods

### Mice

Wild-type C57BL/6J mice of 8-11 weeks age were used in all experiments. Both male and female mice were included in the study. Mice were housed in a semi-barrier animal facility in cages under controlled environmental conditions. The animal room was maintained on a 12-hour light/12-hour dark cycle, with lights on from 07:00 to 19:00, a temperature of 72 ± 2 °F, and relative humidity of 50 ± 10%. Animals had ad libitum access to standard rodent chow and tap water. Cages contained wooden chip bedding and environmental enrichment materials to support animal welfare and natural behaviours.

All animal experiments were conducted in accordance with institutional and national ethical guidelines for animal research and were approved by the relevant animal care and use committees at Nanyang Technological University (Singapore, Animal work license #A25025). The study was designed, conducted, and reported in accordance with the ARRIVE Guidelines 2.0.

### Sample preparation for immunostaining of soft tissues

Freshly dissected murine organs were immediately fixed in 4% paraformaldehyde (Sigma-Aldrich, SDBB6155) for 4 hours at 4°C. Post-fixation organs were immersed overnight in a solution containing 20% sucrose (Bio basic, SB0498) and 2% polyvinylpyrrolidone (PVP; Sigma-Aldrich, P5288) at 4°C followed by incubation in ice- cold solution containing 30% polyethylene glycol and 0.5% Triton X-100 (Sigma- Aldrich, X114). Prior to cryosectioning, organs were embedded and frozen in 8% gelatin (Sigma-Aldrich, G2625) supplemented with 20% sucrose and 2% PVP. Samples were then sectioned at 70-80 μm thickness by a Leica CM3050 cryostat with low-profile blades. Sections were air-dried and stored at -20°C.

### Sample preparation for immunostaining of bones

For preparation of bone tissues, freshly dissected murine bones were immediately fixed in 4% paraformaldehyde for 4 hours at 4°C. Fixed bones were then decalcified with 0.5 M EDTA (Bio-Rad 161-0729) with constant shaking at 4°C for 24 hours. After thorough washing with PBS, decalcified bones were then immersed overnight into 20% sucrose and 2% PVP solution. Finally, bone tissues were embedded and frozen in 8% gelatin in the presence of 20% sucrose and 2% PVP. Samples were then sectioned at 70-80 μm thickness by a Leica CM3050 cryostat with low-profile blades and allowed to air-dry before storing at -20°C.

### Immunostaining

For immunostaining, sections were air-dried for 15 minutes, hydrated with PBS for 5 minutes, permeabilized with 0.3% Triton X-100 for 15 minutes, and blocked in 5% donkey serum for 30 minutes at room temperature (RT). After blocking, sections were incubated with primary antibodies (listed in **Table 2**) diluted in 5% donkey serum for 3 hours at RT. After PBS washing, sections were then incubated with appropriate Alexa Fluor–coupled secondary antibodies (listed in **Table 2)** for 1.5 hours at RT. Finally, sections were washed thoroughly, nuclear counterstaining was performed with 4′,6- diamidino-2-phenylindole (DAPI) and mounting on coverslips using Fluoromount-G (SouthernBiotech 0100-01) was done.

### Image acquisition and analysis

Zeiss LSM880 laser scanning confocal microscope and Leica DM6 B Thunder microscope were used to acquire 3D images of thick immunofluorescent tissue sections. A 20× Plan Apo/0.8 dry lens was employed for acquisition by Zeiss LSM880 microscope which was equipped with seven laser lines (405, 453, 488, 514, 561, 594, and 633 nm), Axio Examiner stand and Colibri 7 epifluorescence light source with LED (light-emitting diode) illumination, fast scanning stage with PIEZO XY, 32-channel gallium arsenide phosphide detector (GaAsP) PMT (photomultiplier tube) plus two- channel standard PMT, acquisition, and analysis software including measurement, multichannel, panorama, manual extended focus, image analysis, time lapse, Z Stack, extended focus, autofocus, and with additional modules: Experiment Designer and Tiles and Position. Thunder Computational Clearing was applied during acquisition with Leica DM6 B Thunder microscope for better contrast and reduced background haze. All the imaging datasets obtained from these microscopes were subsequently processed and reconstructed in three dimensions using Imaris software (version 10.2.0).

### Single cell RNA-sequencing data analysis

The scRNA-seq datasets of interest were obtained from publicly available repositories, including the Gene Expression Omnibus (GEO) and BioStudies databases. Sequencing data for each dataset was imported into R version 4.5.2 for processing using the Seurat R toolkit 5.4.0^69^ individually. Seurat object for each sample was created utilizing the counts of cells with more than 500 unique genes each expressed at least in 3 cells. Then fraction of mitochondrial transcripts was computed for each cell using *PercentageFeatureSet* function with “^mt-” pattern. For downstream analyses, cells with more than 6,000 detected genes, or greater than 25% mitochondrial transcripts were excluded.

Counts of all samples were normalised using *NormalizeData* function with “LogNormalize” method. Top 2,000 variable features for all samples were found using *FindVariableFeatures* function with “variance stabilizing transformation”. Genes annotated to the Gene Ontology term “cell cycle” (GO:0007049) were excluded from the variable feature set prior to data integration. To remove any potential batch effects between samples, integration of all samples was performed using anchors found with *FindIntegrationAnchors* function at default parameters. Integrated data was centered and scaled with *ScaleData* function. Principal components (PCs) were computed by *RunPCA* function and reasonable PCs were selected guided by elbow plots (*ElbowPlot*). Further, non-linear dimensionality reduction was performed with UMAP (*RunUMAP*) and graph-based clustering following “Louvain algorithm” was carried out with *FindNeighbors* and *FindClusters* functions across a range of resolutions. Clusters were manually annotated based on the expression of known canonical markers (**Table 3**) visualized with *FeaturePlot*, *VlnPlot*, and *DotPlot*.

For multi-dataset integration-based endothelial analyses, endothelial cell clusters identified from individual datasets were extracted and merged into a global endothelial cell object, which was processed following the workflow described above. **Table 1** provides list of datasets and specific samples used for the integrated analysis of endothelial cells from healthy murine long bones.

*FindMarkers* function was used to identify genes enriched in each cluster. Genes with an adjusted p-value less than 0.05 and average log_2_(fold change) greater than 0.5 were used for biological interpretation. Gene ontology term signatures were retrieved from Molecular Signatures Database, and per-cell scores were computed using *AddModuleScore* function from Seurat.

RNA velocity analysis was performed using scVelo^70^ (v0.3.3) in Python. Spliced and unspliced transcript count matrices generated by velocyto were matched to the integrated Seurat object using sample identifiers and cell barcodes. Cells successfully matched between the Seurat object and loom files were retained for downstream analyses. The matched AnnData object was preprocessed using scv.pp.filter_and_normalize, followed by neighborhood graph construction and moment estimation using scv.pp.moments. RNA velocities were estimated using the stochastic model implemented in scv.tl.velocity, and transition probabilities were computed using scv.tl.velocity_graph. Velocity stream visualization, velocity confidence, velocity pseudotime, and PAGA graph analyses were performed using the corresponding functions in scVelo. For selected datasets, transcriptional dynamics were recovered using scv.tl.recover_dynamics, and latent time was inferred using scv.tl.latent_time.

### Quantification and statistical analysis

Sample sizes were not pre-determined based on statistical power calculations. Mice were allocated to experiments randomly, and samples were processed in an arbitrary order to minimize potential bias. No blinding was performed during the conduct of the experiments or data analysis, and no animals were excluded from the analysis. Statistical significance of differential gene expression across clusters was tested using Wilcoxon rank-sum test. Genes with p < 0.05 and q < 0.05 were considered to have significant differential expression.

## Legends to Supplementary Figures

**Supplementary Fig 1.** Schematic showing computational and experimental analysis. Integrated single cell and imaging analyses reveal venous and novel fatty acid-handling identity of proposed “type R” endothelial cell population

**Supplementary Fig 2.**
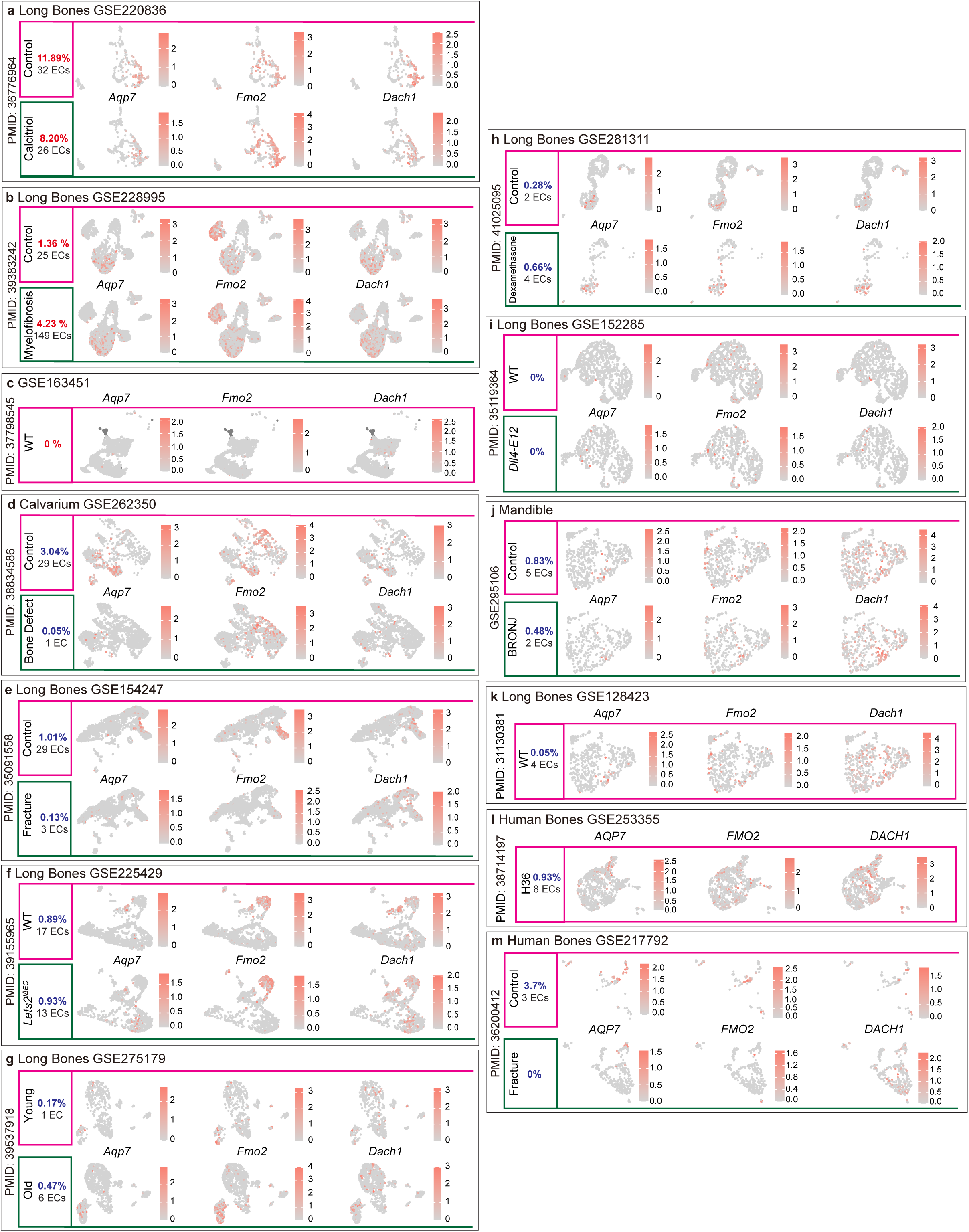
Multi-dataset re-assessment of proposed “type R” endothelial signature. **a**, scRNA-seq dataset, GSE220836, profiling stromal cells from murine long bone under control and calcitriol treatment conditions. Feature plots showing expression of *Aqp7*, *Fmo2* and *Dach1* across entire endothelial cell population. The fraction and number of type R ECs identified in each condition are indicated. **b**, scRNA-seq dataset, GSE228995, profiling stromal cells from murine long bone from control and myelofibrotic mice. Feature plots showing expression of *Aqp7*, *Fmo2* and *Dach1* across entire endothelial cell population. The fraction and number of type R ECs identified in each condition are indicated. **c**, scRNA-seq dataset, GSE163451, profiling stromal cells from murine long bone from young WT mice. Feature plots showing expression of *Aqp7*, *Fmo2* and *Dach1* across entire endothelial cell population. The fraction and number of type R ECs identified in each condition are indicated. **d**, scRNA-seq dataset, GSE262350, profiling stromal cells from murine intact calvarium and calvarium with bone defect (post-injury day 14). Feature plots showing expression of *Aqp7*, *Fmo2* and *Dach1* across entire endothelial cell population. The fraction and number of type R ECs identified in each condition are indicated. **e**, scRNA-seq dataset, GSE154247, profiling stromal cells from murine intact and fractured long bone (post-fracture day 14). Feature plots showing expression of *Aqp7*, *Fmo2* and *Dach1* across entire endothelial cell population. The fraction and number of type R ECs identified in each condition are indicated. **f**, scRNA-seq dataset, GSE225429, profiling stromal cells from WT and *Lats2^iΔEC^*. Feature plots showing expression of *Aqp7*, *Fmo2* and *Dach1* across entire endothelial cell population. The fraction and number of type R ECs identified in each condition are indicated. **g**, scRNA-seq dataset, GSE275179, profiling stromal cells from WT young and old murine long bones. Feature plots showing expression of *Aqp7*, *Fmo2* and *Dach1* across entire endothelial cell population. The fraction and number of type R ECs identified in each condition are indicated. **h**, scRNA-seq dataset, GSE281311, profiling combined stromal and hematopoietic cells from the long bone under control and dexamethasone treatment conditions. Feature plots showing the expression of *Aqp7, Fmo2*, and *Dach1* across entire endothelial cell population. The fraction and number of type R ECs identified in each condition are indicated. **i**, scRNA-seq dataset, GSE152285, profiling stromal cells from long bones of wild-type (WT) and *Dll4-E12* mice. Feature plots showing the expression of *Aqp7, Fmo2,* and *Dach1* across entire endothelial cell population. The fraction and number of type R ECs identified in each condition are indicated. **j**, scRNA-seq dataset, GSE295106, profiling stromal cells from mandible bone under control and BRONJ (Bisphosphonate-Related Osteonecrosis of the Jaw) conditions. Feature plots showing the expression of *Aqp7, Fmo2,* and *Dach1* across entire endothelial cell. The fraction and number of type R ECs identified in each condition are indicated. **k**, scRNA-seq dataset, GSE128423, profiling stromal cells from long bones of young WT mice. Feature plots showing the expression of *Aqp7*, *Fmo2*, and *Dach1* across endothelial cell populations. The fraction and number of type R ECs identified are indicated. **l**, scRNA-seq dataset, GSE253355, profiling enriched stromal and hematopoietic stem/progenitor cells from human bone samples. Feature plots showing *AQP7, FMO2,* and *DACH1* expression across entire endothelial cell population of H36 sample. The fraction and number of type R ECs are indicated. **m**, scRNA-seq dataset, GSE217792, profiling cells of human intramedullary canal tissue under control and fracture conditions. Feature plots showing *AQP7, FMO2,* and *DACH1* expression across endothelial cell populations. The fraction and number of type R ECs are indicated for each condition.

**Supplementary Fig 3.** Re-clustering analyses shows consistent incomplete co-expression of “type R” markers. **a**, Schematic illustrating differences in marker selection between single-cell transcriptomics (*Aqp7*, *Fmo2*) and imaging-based analyses (Emcn, Cav1) of type R Feature plots showing expression of *Aqp7*, *Fmo2*, *Dach1*, *Vwf* and *Selp* in total ECs. Arterial ECs were defined as *Cav1^+^ Bmx^+^ Sox17^+^* cells. The fraction and number of type R and arterial ECs is indicated. Violin plots showing expression of *Emcn, Cav1* and *Flt4* across EC sub-populations. Table showing fraction of cells in original “type R” cluster expressing *Aqp7* and *Fmo2* (taken from PMID: 39528700). **b**, scRNA-seq dataset, GSE154247, profiling stromal cells from murine intact and fractured long bone (post-fracture day 14). Feature plots showing expression of *Aqp7, Fmo2* and *Dach1* across entire endothelial cell population. The fraction and number of type R ECs identified in each condition are indicated. Violin plots showing expression of *Emcn, Cav1* and *Flt4* across EC sub-populations. scRNA-seq dataset, GSE275179, profiling stromal cells from WT young and old murine long bones. Feature plots showing expression of *Aqp7, Fmo2* and *Dach1* across entire endothelial cell population. The fraction and number of type R ECs identified in each condition are indicated. Violin plots showing expression of *Emcn, Cav1* and *Flt4* across EC sub-populations. **c**, UMAP showing type R ECs identified in integrated analysis of endothelial cells from 10 healthy murine long bone scRNA-seq datasets. The fraction and number of type R ECs is indicated. Note two datasets were found to have no type R ECs.

**Supplementary Fig 4.** Single cell transcriptomic and imaging-based reassessment of DACH1 in bone and marrow. **a**, scRNA-seq dataset, GSE156636, profiling stromal cells from long bones of juvenile (P21) mice. UMAP showing capillary and venous EC populations in juvenile murine long bone ECs. Violin plots showing expression of *Aqp7, Fmo2, Dach1, Ptgs1, Vwf, Lrg1, Selp,* and *Nr2f2* in capillary vs venous ECs. The fraction and number of type R ECs is indicated. **b**, UMAPs showing capillary and venous endothelial cells (ECs) in control and *Dach1* overexpressing (*Dach1^OE^*) murine long bones from GSE239627 dataset. Violin plots showing expression of *Aqp7, Fmo2, Vwf, Lrg1, Selp, Nr2f2, Ptgs1,* and *Dach1* in capillary vs venous ECs in both conditions. **c**, UMAP showing various cell types in control and *Dach1^OE^* groups. Feature plots showing expression of *Aqp7, Fmo2, Dach1, C1qtnf9, Emcn,* and *Cav1* in control vs *Dach1^OE^* groups. The fraction and number of type R ECs in both groups is indicated. Bar plot depicting in *Dach1*^+^ cell fraction in control vs *Dach1^OE^* mice. **d**, Left panel: Representative images of metaphyseal (MP) region of young murine tibia showing immunolabelling for DACH1 (green) and Endomucin (red) and nuclear staining with DAPI (blue). Right panel: Representative images of MP and diaphyseal (DP) regions of young murine tibia showing immunolabelling for DACH1 (green), RUNX2 (red) and Osteopontin (white) and nuclear staining with DAPI (blue). Scale bars: 50 μm (white) and 15 μm (orange). **e**, Representative images of transition zone (TZ) of young murine tibia showing immunolabelling for DACH1 (green) and Endomucin (red) and nuclear staining with DAPI (blue). Scale bars: 50 μm (white)

**Supplementary Fig 5.** Endothelial cluster separation is subject to input resolution. **a**, Impact of resolution on clustering on integrated data of 10 healthy murine long bone scRNA-seq datasets. UMAPs showing endothelial clustering at low and high resolution values. **b,** Feature plots showing expression pattern of *Fmo2* and *Aqp7* in entire endothelial population. **c,** UMAP showing various endothelial sub-populations in dataset GSE239627. Violin plots showing expression of *Lpl*, *Tcf15*, *Meox2*, *Cav1*, *Cd36* and *Rgcc* across endothelial sub-populations.

**Supplementary Fig 6.** Fatty acid-handling endothelial cells in craniofacial skeleton. **a**, scRNA-seq dataset, GSE262350, profiling stromal cells from young murine calvarium. UMAP showing various endothelial sub-populations in murine calvarium. Violin plots showing expression of *Lpl*, *Tcf15*, *Meox2*, *Cav1*, *Cd36* and *Rgcc* across endothelial sub-populations. **b**, scRNA-seq dataset, GSE295106, profiling mandibular stromal cells from young mice. UMAP showing various endothelial sub-populations in murine mandible. Violin plots showing expression of *Lpl*, *Tcf15*, *Meox2*, *Cav1*, *Cd36* and *Rgcc* across endothelial sub-populations. **c**, scRNA-seq dataset, GSE316924, profiling mandibular stromal cells from young mice. UMAP showing various endothelial sub-populations in murine mandible. Violin plots showing expression of *Lpl*, *Tcf15*, *Meox2*, *Cav1*, *Cd36* and *Rgcc* across endothelial sub-populations. **d**, Left panel: Representative images of young murine mandible showing immunolabelling for RGCC (green), Endomucin (red) and VEGFR3 (white) and nuclear staining with DAPI (blue). Right panel: Representative images of young murine mandible showing immunolabelling for RGCC (green), Endomucin (red) and VEGFR3 (cyan) and nuclear staining with DAPI (blue). Scale bars: 50 μm (white) and 15 μm (magenta).

